# 5-Deazaflavin (TND1128) and its hybrid analogs are cytoprotective against hydrogen peroxide (H_2_O_2_)-induced oxidative stress

**DOI:** 10.1101/2024.05.07.592882

**Authors:** Kaori Kubota, Shutaro Katsurabayashi, Takuya Watanabe, Katsunori Iwasaki, Tomohisa Nagamatsu, Norio Akaike

**Author notes:** Correspondence: Dr. Kaori Kubota, Department of Neuropharmacology, Faculty of Pharmaceutical Sciences, Fukuoka University, 8-19-1 Nanakuma, Jonan-ku, Fukuoka 814-0180, Japan. Department of Pharmaceutical Sciences, International University of Health and Welfare, 137-1 Enokizu, Okawa City, Fukuoka 831-8501, Japan (Kaori Kubota). **List of e-mail addresses and ORCID numbers of authors:** Kaori Kubota Shutaro Katsurabayashi Takuya Watanabe Katsunori Iwasaki Tomohisa Nagamatsu Norio Akaike.

## Abstract

Increased production of reactive oxygen species (ROS) and oxidative stress are implicated in mitochondrial dysfunction, contributing to the pathogenesis of many neurodegenerative diseases. Research is ongoing into a new treatment approach for neurodegeneration, focusing on reactivating dysfunctional mitochondria. Some 5-deazaflavins, such as 10-ethyl-3-methylpyrimido[4,5-*b*]quinoline-2,4(3*H*,10*H*)-dione (TND1128), and four analogs of 5-deazaflavin, including β-nicotinamide mononucleotide (β-NMN), demonstrate efficient self-redox abilities similar to β-NMN, making them potential activators of mitochondrial energy synthesis. This study examines whether TND1128 and its analogs have protective effects against cellular impairment induced by oxidative stress. These compounds exhibit proliferative potential against normal cells. Moreover, TND1128 and its analogs significantly improved cell viability against hydrogen peroxide (H_2_O_2_)-induced oxidative stress injury. Our study confirms the cytoprotective effect of these 5-deazaflavins through mitochondrial activation. We anticipate TND1128 and its analogs will serve as mitochondria-stimulating drugs capable of rescuing deteriorating neurons in aging or diseases.

## 1. Introduction

Mitochondria, which are present in almost all cells, are vital organelles involved not only in energy production but also in regulating Ca^2+^ concentration and reducing oxidative stress. Recently, it has been discovered that mitochondrial dysfunction contributes to chronic inflammation and is associated with aging and the onset of various diseases such as cancer, dementia, mental illnesses like depression, and diabetes. Research is ongoing on a new treatment method aimed at extending the healthy lifespan of humans by reactivating dysfunctional mitochondria.

β-Nicotinamide mononucleotide (β-NMN) is a precursor of the coenzyme nicotinamide adenine dinucleotide (NAD^+^), essential for cell activity, and is present in the cells of all living organisms (Fig. 1a). Typically, β-NMN is synthesized in the body from materials such as vitamin B3, but its production decreases with aging. Consequently, NAD^+^ production diminishes, leading to further damage to the cell nucleus and decreased mitochondrial activity. β-NMN is suggested to have neuroprotective and anti-aging effects due to the production of NAD^+^ from β-NMN. NAD^+^ serves as a cofactor for various processes within mitochondria, sirtuins, known as longevity genes, and poly [ADP-ribose] polymerase involved in DNA repair (1, 2). Additionally, β-NMN has beneficial effects on the central nervous system (CNS), such as enhancing mitochondrial function in the brain (3), preventing β-amyloid oligomer-induced cognitive impairment (4), and delaying astrocyte-mediated motor neuron death (5). β-NMN is found in foods such as green and yellow vegetables and fruits, but the amount contained is often insufficient. Given the challenge of obtaining sufficient amounts from food, there is a need for alternative drugs or supplements that can be taken safely and easily.

**Figure 1.**
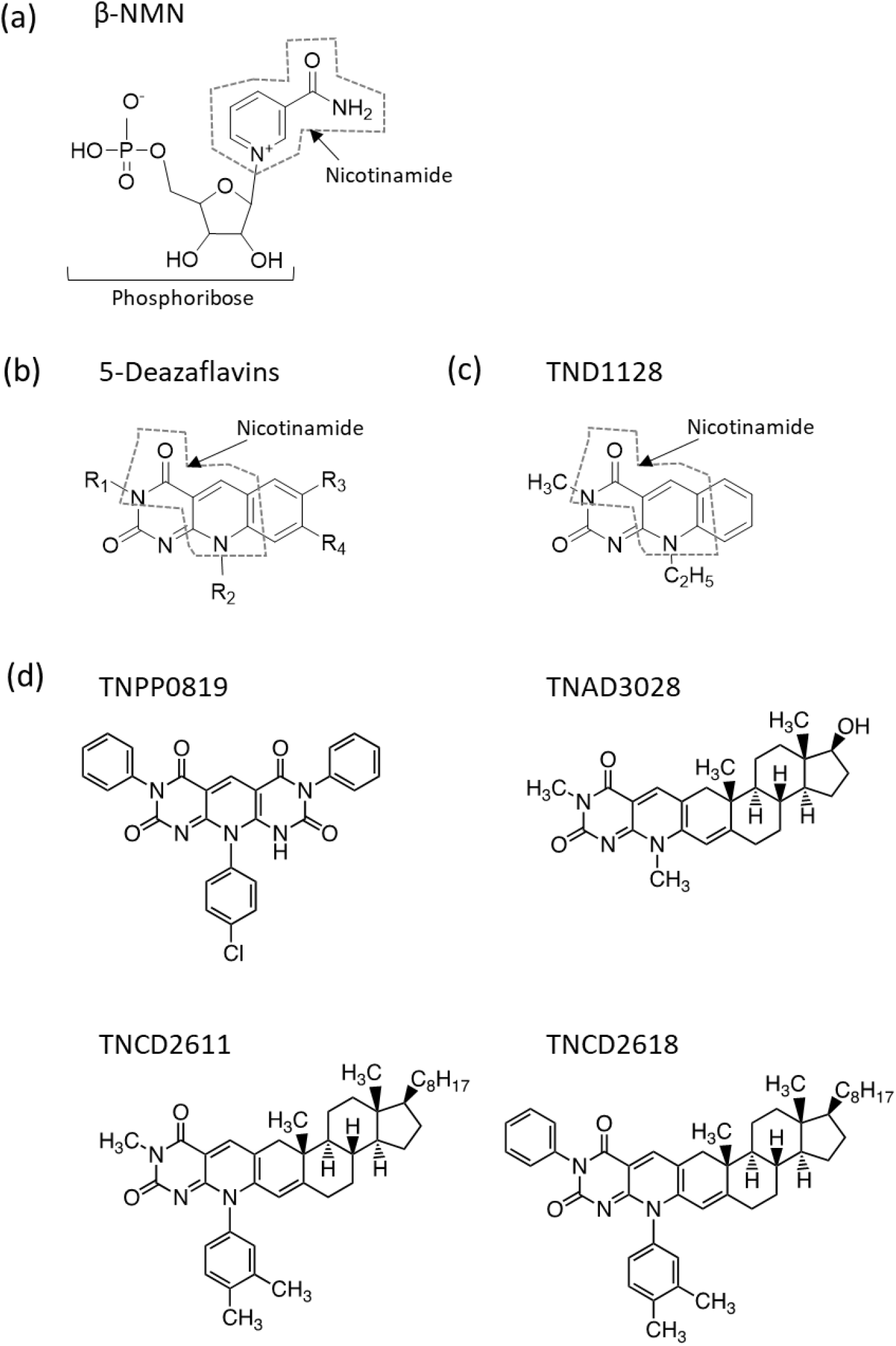
Structures of β-NMN, TND1128, and other 5-deazaflavin analogs. (a) β-NMN: a highly water-soluble compound with a phosphoribosyl bond at the N-position of nicotinamide. This compound is known for its effective redox activity. (b) Basic structure of 5-deazaflavin. (c) **TND1128**: a hydrophobic compound synthesized as a 5-deazaflavin derivative. This compound also contains a nicotinamide structure. (d) Structures of four 5-deazaflavin analogs. Each compound has the following structural formula: pyridodipyrimidine (**TNPP0819**), testosterone-deazaflavin hybrid (**TNAD3028**), and cholesterol-deazaflavin hybrids (**TNCD2611** and **TNCD2618**).

Extensive research has been conducted on the synthesis of a compound possessing a flavin skeleton and pharmacological effects. Derivatives of 5-deazaflavin (pyrimido[4,5*-b*]quinoline-2,4(3*H*,10*H*)-diones) feature a unique chemical structure obtained by substituting N at position 5 of riboflavin with CH (Fig. 1b) and exhibit redox functions similar to those of NAD^+^. These derivatives also activate intracellular ATP production and sirtuin gene function. TND1128, a representative 5-deazaflavin (10-ethyl-3-methylpyrimido[4,5-*b*]quinoline-2,4(3*H*,10*H*)-dione), is known for its efficient self-redox ability (6, 7). Furthermore, TND1128 has been reported to enhance ATP production in cultured astrocytes (8), promote the elongation and branching of dendrites in cultured neurons (9), and exert protective effects in the mouse brain (10). However, few studies have investigated the neuroprotective effects of 5-deazaflavins other than TND1128. With the aim of application in the treatment of neuropsychiatric diseases such as Alzheimer’s disease and depression, various 5-deazaflavin-derived and analogous compounds with increased lipophilicity have been synthesized to enhance their transit to the central nervous system (8). Through screening the neurite outgrowth effects of seven of these derivatives, four analogs such as pyridodipyrimidine (TNPP0819) (11), testosterone-deazaflavin hybrid (TNAD3028) (12), and cholesterol-deazaflavin hybrids (TNCD2611 and TNCD2618) (13) (Fig. 1d) were identified that significantly promote the elongation and branching of dendrites in cultured neurons (8).

In the present study, we examined the protective effect of TND1128 and its analogs against hydrogen peroxide (H_2_O_2_)-induced oxidative stress to evaluate their potential as activators of energy production and compared their efficacy with that of β-NMN as a control.

## 2. Materials and Methods

### 2.1. Materials

TND1128 (10-ethyl-3-methylpyrimido[4,5-*b*]quinoline-2,4(3*H*,10*H*)-dione) (10-ethyl-3-methyl-5-deazaflavin) and four analogs, **TNPP0819** (10-(4-chlorophenyl)-3,7-diphenylpyrido[2,3-*d*:6,5-*d*’]dipyrimidine-2,4,6,8(1*H*,3*H*,7*H*,10*H*)-tetraone), **TNAD3028** (5’-deaza-17b-hydroxy-3’,8’-dimethylandrost-2,4-dieno[2,3-*g*]pteridine-2’,4’(3’*H*,8’*H*)-dione), **TNCD2611** (3’-methyl-8’-(3,4-dimethylphenyl)-5’-deazacholest-2,4-dieno[2,3-*g*]pteridine-2’,4’(3’*H*,8’*H*)-dione), and **TNCD2618** (8’-(3,4-dimethylphenyl)-3’-phenyl-5’-deazacholest-2,4-dieno[2,3-*g*]pteridine-2’,4’(3’*H*,8’*H*)-dione), synthesized by the co-author (T.N.) (Fig. 1), were employed in the study (8). β-NMN and dimethylsulfoxide (DMSO) were procured from Sigma-Aldrich (St. Louis, USA). Hydrogen peroxide (H_2_O_2_) was procured from FUJIFILM Wako (Osaka, Japan).

### 2.2. Cell cultures

Rat pheochromocytoma (PC12) cells were obtained from the American Type Culture Collection (Manassas, Virginia, USA). These cells were cultured in RPMI-1640 medium (Nissui, Tokyo, Japan) supplemented with 5% (v/v) fetal bovine serum (Thermo Fisher Scientific, Waltham, MA, USA), 10% (v/v) horse serum (Thermo Fisher Scientific, Waltham, MA, USA), 100 units/ml penicillin, and 100 μg/ml streptomycin (FUJIFILM Wako, Osaka, Japan). The cells were grown until confluence at 37°C in a 5% CO_2_ atmosphere. The medium was refreshed two or three times per week.

### 2.3. Cell viability assay

The effect of TND1128 and its analogs on the cell viability of PC12 and H_2_O_2_-injured PC12 cells was determined using a colorimetric assay with the Cell Counting Kit-8 (CCK-8, Dojindo, Japan) following the manufacturer’s instructions. Briefly, PC12 cells were seeded in 96-well plates at a density of 2 × 10^4^ cells/well for 24 h. Subsequently, the cells were treated with increasing concentrations of β-NMN, TND1128, and its derivatives (0.01–1.0 μM, dissolved in DMSO) or vehicle (DMSO) for 24 h. CCK-8 reagent was added, and the absorbance of the resulting formazan at 450 nm was measured using an automatic multi-well spectrophotometer (Tecan, Männedorf, Switzerland). The absorbance values for each treatment group were expressed as a percentage of the control.

Furthermore, the effect of β-NMN, TND1128, and its analogs on the reduction of H_2_O_2_-induced cytotoxicity was evaluated. After seeding PC12 cells in 96-well plates at a concentration of 2 × 10^4^ cells/well for 24 h, the cells were treated with β-NMN, TND1128, and its derivatives for 2 h prior to H_2_O_2_ exposure. Subsequently, 20 μM of H_2_O_2_ or β-NMN, TND1128, and its analogs were added to the culture medium and incubated for 24 h. To assess cell viability, the H_2_O_2_-containing media were discarded, and cell viability was determined using the CCK-8 assay.

### 2.4. Statistical analysis

All statistical analyses were conducted using GraphPad Prism 8 software (La Jolla, CA, USA). The results were expressed as the mean ± standard error of the mean (SEM). For multiple comparisons between the treated conditions and control, one-way ANOVA with post hoc Tukey’s multiple comparisons tests or Dunnett’s multiple comparisons tests were performed. The criterion for statistical significance was set at p<0.05.

## 3. Results

### 3.1. Effects of TND1128 and 5-deazaflavin derivatives on cell proliferation

We examined the effects of **TND1128**, a representative 5-deazaflavin, and **β-NMN** on the cell proliferation of normal cells (Fig. 2). Treatment of PC12 cells with **TND1128** for 24 h resulted in a significant activation of cell proliferation within a concentration range of 0.01–1.0 μM (F (5, 158)=8.679, p=0.0193 for 0.01 μM, p=0.0002 for 0.03 μM, p=0.0036 for 0.1 μM, p=0.0071 for 0.3 μM, p<0.0001 for 1.0 μM vs. Control; Fig. 2a). In contrast, β-NMN, widely recognized for its NAD^+^-like effects, demonstrated significant cell proliferation within the concentration range of 0.1–1.0 μM (F (5, 158)=3.883, p=0.0432 for 0.1 μM, p=0.0099 for 0.3 μM, p=0.0197 for 1.0 μM vs. Control; Fig. 2b). These results suggest that **TND1128** exhibits a strong NAD^+^-like effect even at lower concentrations.

**Figure 2.**
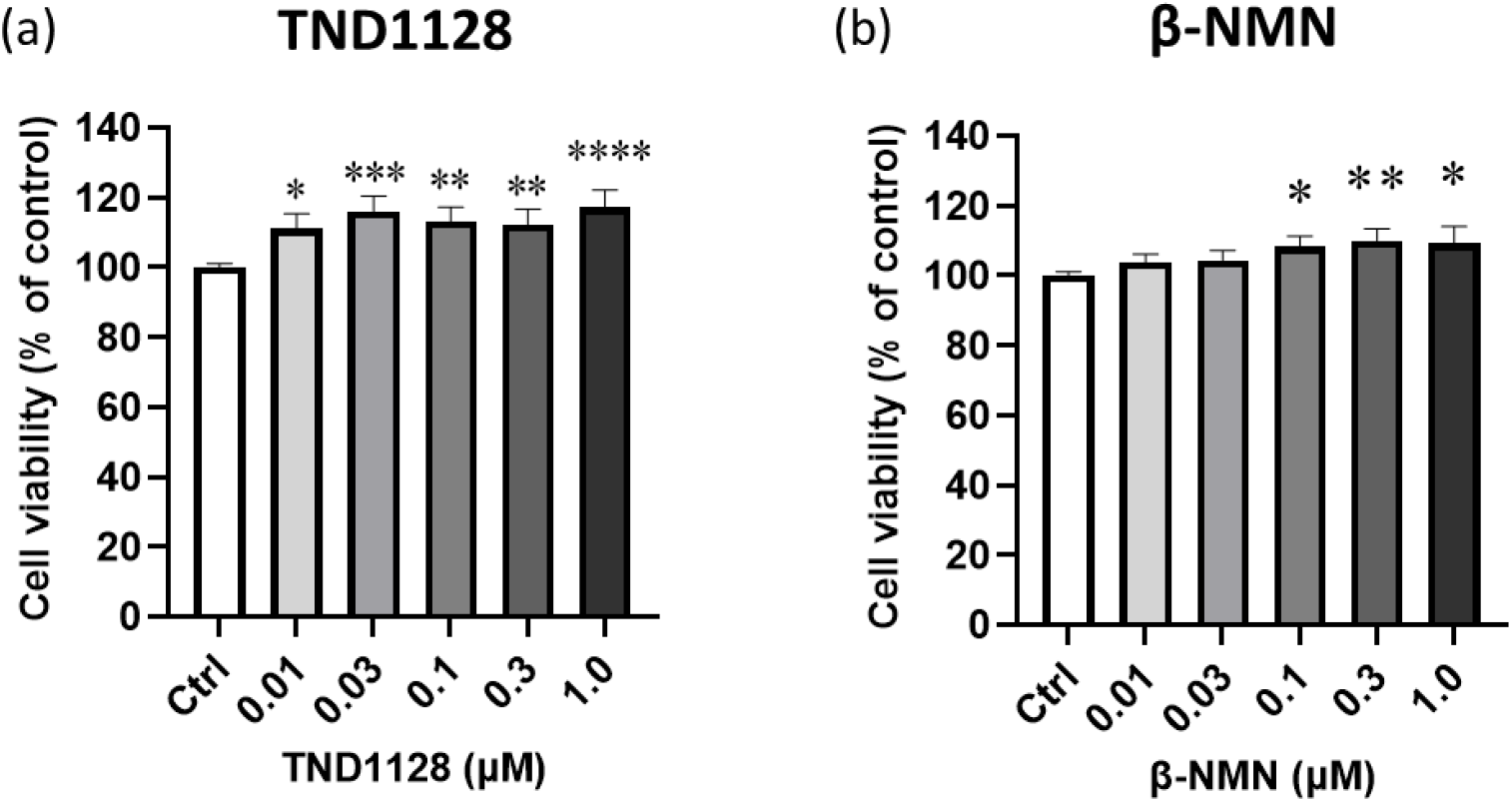
Influence of TND1128 and β-NMN on cell proliferation. PC12 cells were treated with the indicated doses of TND1128 (a, 0–1.0 µM) or β-NMN (b, 0–1.0 µM) for 24 h. Cell proliferation was assessed using the CCK-8 assay and expressed as a percentage of control. *p<0.05, **p<0.01, ***p<0.001, and ****p<0.0001 indicate significant differences compared to the Ctrl group (ANOVA followed by Dunnett’s multiple comparisons test).

Next, we investigated the effects of 5-deazaflavin compounds other than TND1128 on cell proliferation (Fig. 3). When PC12 cells were treated with each compound for 24 h, all compounds significantly increased cell proliferation (**TNPP0819**: F (5, 158)=18.55, p=0.0018 for 0.01 μM, p<0.0001 for 0.03 μM, p<0.0001 for 0.1 μM, p<0.0001 for 0.3 μM, p<0.0001 for 1.0 μM vs. Control; Fig. 3a, **TNAD3028**: F (5, 158)=7.800, p=0.0053 for 0.03 μM, p=0.0100 for 0.1 μM, p=0.0015 for 0.3 μM, p<0.0001 for 1.0 μM vs. Control; Fig. 3b, **TNCD2611**: F (5, 158)=6.961, p=0.0041 for 0.01 μM, p=0.0097 for 0.03 μM, p=0.0018 for 0.1 μM, p=0.0008 for 0.3 μM vs. Control; Fig. 3c, **TNCD2618**: F (5, 158)=7.820, p=0.0479 for 0.01 μM, p=0.0251 for 0.03 μM, p<0.0001 for 0.1 μM, p=0.0002 for 0.3 μM vs. Control; Fig. 3d). Among the four compounds examined in this study, **TNPP0819** appeared to be particularly potent compared to the positive control β-NMN (Fig. 3a). These results suggest that 5-deazaflavin compounds have a significant impact on cell proliferation. Furthermore, no significant cytotoxicity was observed for any of the compounds at the concentrations treated in this study.

**Figure 3.**
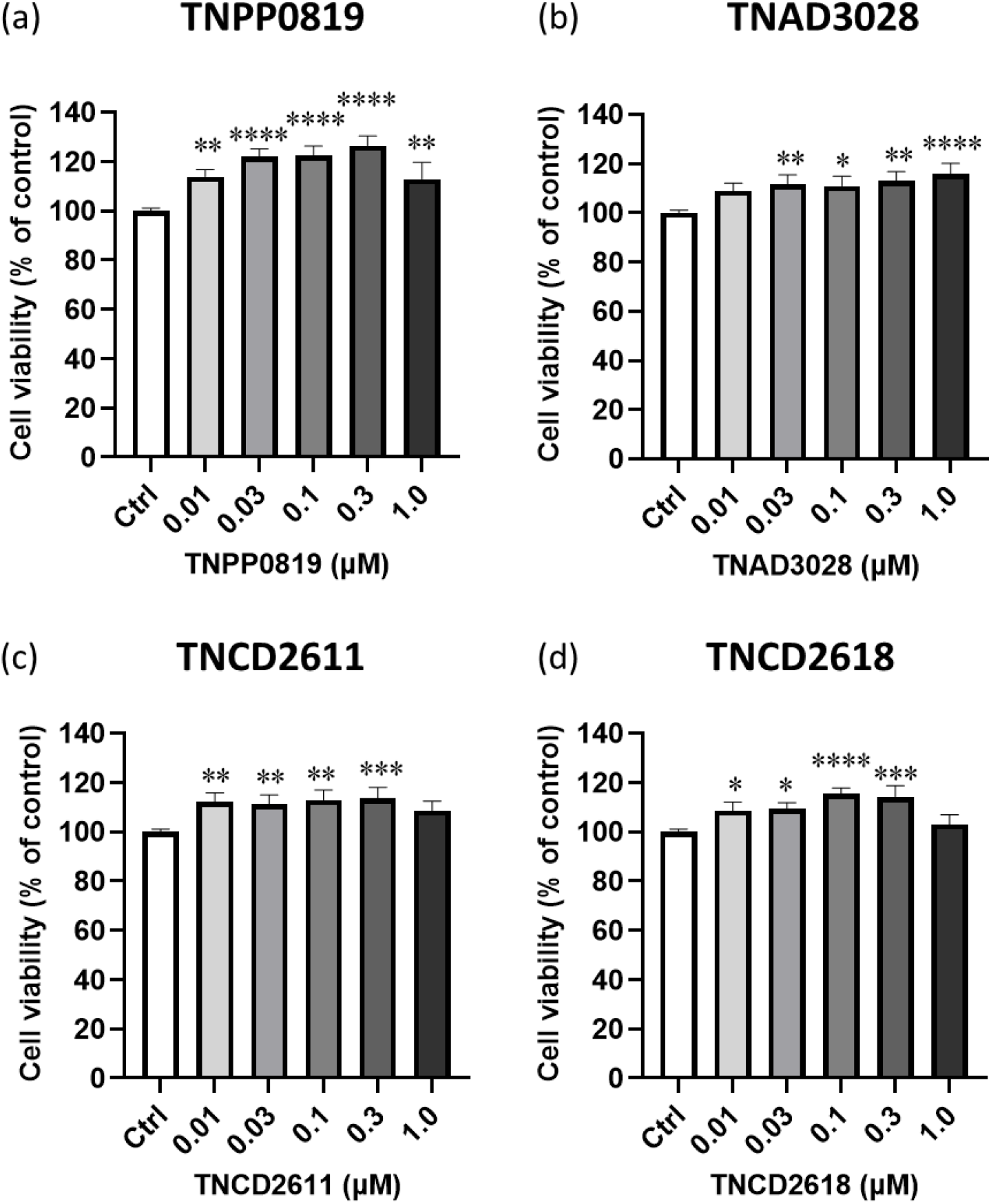
Influence of other 5-deazaflavin analogs on cell proliferation. PC12 cells were treated with the indicated doses of 5-deazaflavin analogs (0–1.0 µM, a: **TNPP0819**, b: **TNAD3028**, c: **TNCD2611**, d: **TNCD2618**) for 24 h. Cell proliferation was assessed using the CCK-8 assay and expressed as a percentage of the control. *p<0.05, **p<0.01, ***p<0.001, and ****p<0.0001 indicate significant differences compared to the Ctrl group (ANOVA followed by Dunnett’s multiple comparisons test).

### 3.2. Protective effects of TND1128 and its analogs against oxidative stress-induced cell death

To examine the effect of 5-deazaflavin compounds on oxidative stress damage, we determined the conditions for inducing cell damage (Fig. 4a). Cell viability was significantly decreased at concentrations exceeding 20 µM of H_2_O_2_ treatment (F (9, 238)=156.3, p<0.0001 for 20–100 μM H_2_O_2_ vs. Control; Fig. 4a). Because an approximately 50% reduction in viability was observed with 20–25 μM treatment, subsequent experiments were conducted at this concentration to induce cell damage.

**Figure 4.**
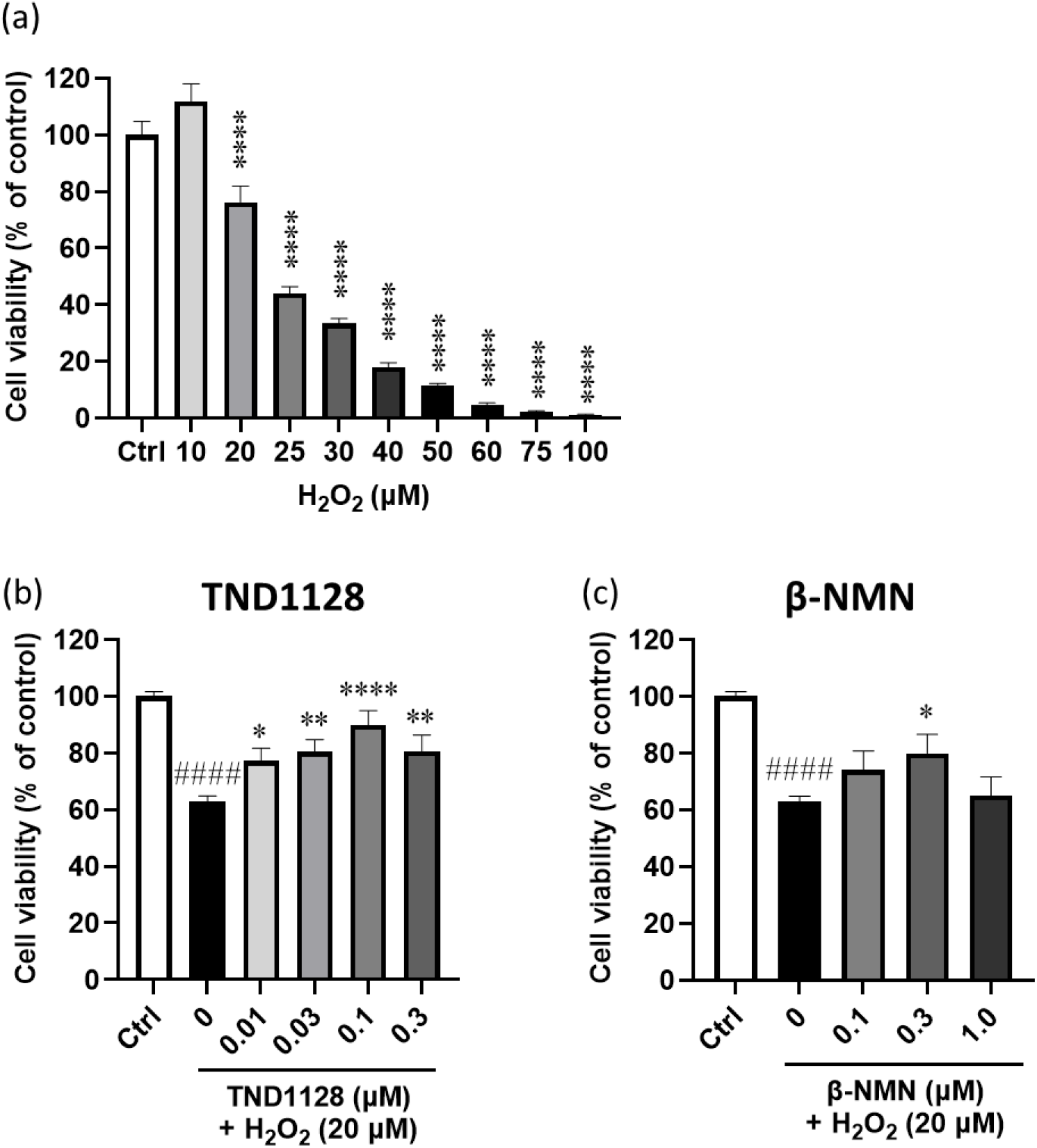
Protective effects of TND1128 and β-NMN on cell viability following H_2_O_2_ exposure. (a) Influence of H_2_O_2_ on cell viability. PC12 cells were treated with the indicated doses of H_2_O_2_ (0–100 µM) for 24 h. Cell viability was assessed using the CCK-8 assay and expressed as a percentage of the control. ****p<0.0001 indicates significant differences compared to the Ctrl group (ANOVA followed by Dunnett’s multiple comparisons test). Exposure to H_2_O_2_≥20 µM significantly induced cytotoxicity. (b, c) PC12 cells were pre-treated with TND1128 (b, 0.01–0.3 µM) or β-NMN (c, 0.1–1.0 µM) for 2 h prior to H_2_O_2_ exposure. After 2 h, 20 μM of H_2_O_2_ with the indicated doses of TND1128 or β-NMN were added to the culture medium and incubated for 24 h. Cell viability was assessed using the CCK-8 assay and expressed as a percentage of the control. ^# # #^p<0.0001 indicates significant differences compared to the Ctrl group; *p<0.05, **p<0.01, and ****p<0.0001 indicate significant differences compared to the 20 μM H_2_O_2_ (+ 0.05% DMSO) group (ANOVA followed by Tukey’s multiple comparisons test).

Next, the protective effects of TND1128 and four 5-deazaflavin analogs on oxidative stress injury of PC12 cells were examined, compared to the positive control β-NMN. Single treatment results indicated that **β-NMN** affected cell proliferation at concentrations of 0.1–1.0 mg/ml, whereas other compounds, including **TND1128**, showed effects at even lower concentrations (Fig. 2, 3). Based on these data, the cytoprotective effect of β-NMN was examined up to a concentration of 1.0 mg/ml, whereas the effects of other compounds were assessed up to a lower dose of 0.3 mg/ml. **TND1128** demonstrated cytoprotective effects, ameliorating the reduction in cell viability induced by H_2_O_2_ treatment in a bell-shaped, dose-dependent manner (F (5, 708)=40.26, p<0.0001 for 20 μM H_2_O_2_ (+ 0.05% DMSO) vs. Control, p=0.0313 for 0.01 μM TND1128 + 20 μM H_2_O_2_, p=0.0034 for 0.03 μM TND1128 + 20 μM H_2_O_2_, p<0.0001 for 0.1 μM TND1128 + 20 μM H_2_O_2_, p=0.0029 for 0.3 μM TND1128 + 20 μM H_2_O_2_ vs. 20 μM H_2_O_2_ (+ 0.05% DMSO); Fig. 4b). Similarly, **β-NMN** also exhibited cytoprotective effects similar to those of TND1128 (F (4, 613)=51.91, p<0.0001 for 20 μM H_2_O_2_ (+ 0.05% DMSO) vs. Control, p=0.0223 for 0.3 μM β-NMN + 20 μM H_2_O_2_ vs. 20 μM H_2_O_2_ (+ 0.05% DMSO); Fig. 4c). However, consistent with the findings in Fig. 3, TND1128 appeared to have a stronger protective effect than β-NMN.

Moreover, we investigated four other 5-deazaflavin analogs, and all exhibited a significant improvement in cell viability (**TNPP0819**: F (5, 720)=39.83, p<0.0001 for 20 μM H_2_O_2_ (+ 0.05% DMSO) vs. Control, p=0.0018 for 0.1 μM TNPP0819 + 20 μM H_2_O_2_, p=0.0073 for 0.3 μM TNPP0819 + 20 μM H_2_O_2_ vs. 20 μM H_2_O_2_ (+ 0.05% DMSO); Fig. 5a, **TNAD3028**: F (5, 720)=38.86, p<0.0001 for 20 μM H_2_O_2_ (+ 0.05% DMSO) vs. Control, p<0.0001 for 0.1 μM TNAD3028 + 20 μM H_2_O_2_, p=0.0177 for 0.3 μM TNAD3028 + 20 μM H_2_O_2_ vs. 20 μM H_2_O_2_ (+ 0.05% DMSO); Fig. 5b, **TNCD2611**: F (5, 720)=45.25, p<0.0001 for 20 μM H_2_O_2_ (+ 0.05% DMSO) vs. Control, p=0.0095 for 0.1 μM TNCD2611 + 20 μM H_2_O_2_; Fig. 5c, **TNCD2618**: F (5, 720)=42.58, p<0.0001 for 20 μM H_2_O_2_ (+ 0.05% DMSO) vs. Control, p=0.0009 for 0.1 μM TNCD2618 + 20 μM H_2_O_2_; Fig. 5d). These four 5-deazaflavin analogs examined in this study demonstrated a cell protective effect similar to that of TND1128 and β-NMN.

**Figure 5.**
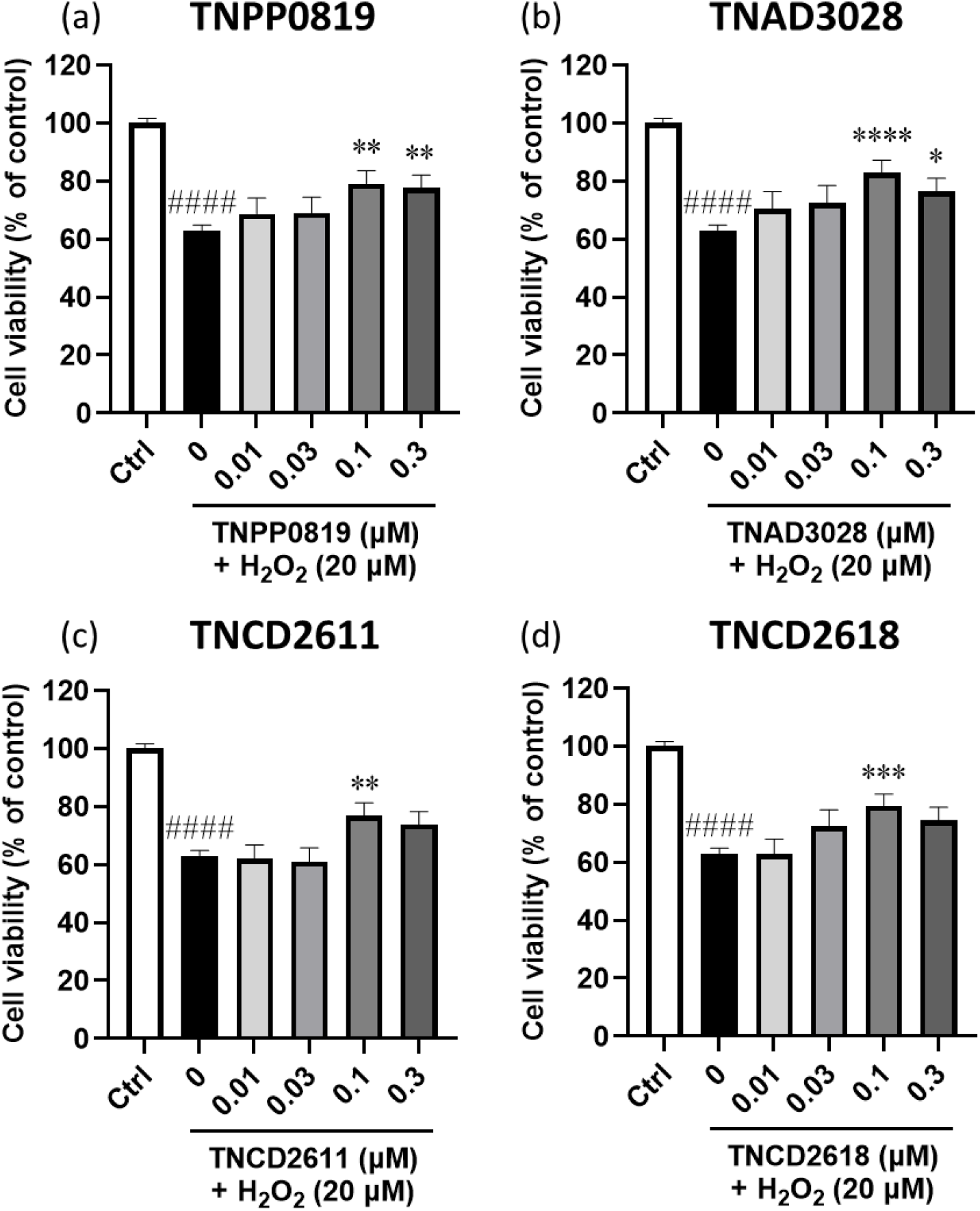
Protective effects of other 5-deazaflavin analogs on cell viability following H_2_O_2_ exposure. PC12 cells were pre-treated with the indicated doses of 5-deazaflavin analogs (0–0.3 µM, a: **TNPP0819**, b: **TNAD3028**, c: **TNCD2611**, d: **TNCD2618**) for 2 h prior to H_2_O_2_ exposure. After 2 h, 20 μM of H_2_O_2_ with the indicated doses of 5-deazaflavin analogs were added to the culture medium and incubated for 24 h. Cell viability was assessed using the CCK-8 assay and expressed as a percentage of control. # # #p<0.0001 indicates significant differences compared to the Ctrl group; *p<0.05, **p<0.01, and ***p<0.001 indicate significant differences compared to the 20 μM H_2_O_2_ (+ 0.05% DMSO) group (ANOVA followed by Tukey’s multiple comparisons test).

## 4. Discussion

Reactive oxygen species (ROS) induce oxidative stress, resulting in damage to organelles and macromolecules such as DNA, proteins, and lipids, ultimately leading to cell death (14). In addition to exogenous ROS inducers, endogenous substances such as 15-deoxy-Δ12,14-prostaglandin J2 (15d-PGJ2), synthesized via prostaglandin H2, also trigger inflammatory responses as endogenous electrophiles (15). The increased production of ROS and oxidative stress contribute to the dysfunction of mitochondrial energy production and have been implicated in the pathogenesis of numerous neurodegenerative diseases, including Alzheimer’s disease (16), Parkinson’s disease (17), and Huntington’s disease (18). Particularly in brain cells with high energy demands, mitochondria are notably susceptible to malfunction due to age-related abnormalities in mitochondrial electron transport.

We have studied a wide range of 5-deazaflavins with increased lipophilicity to improve their translocation into the brain (8). Previous studies have reported that TND1128 acts as a coenzyme factor for activating ATP production in cultured astrocytes (8), significantly reducing cytoplasmic and mitochondrial Ca^2+^ elevation induced by electrical stimulation (10) and promoting significant branching of axons and dendrites in cultured neuronal cells (9). These mechanisms of TND1128 appear analogous to those of β-NMN. In this study, we investigated their cytoprotective effects against oxidative stress using respective compounds characterized by structural formulas representing 5-deazaflavins (TND1128) and four analogs: pyridodipyrimidine (TNPP0819), testosterone-deazaflavin hybrid (TNAD3028), and cholesterol-deazaflavin hybrid (TNCD2611 and TNCD2618) (Fig. 1c and d). TND1128 and the other 5-deazaflavin analogs exhibited both proliferative and cytoprotective effects on PC12 cells similar to β-NMN. Although they ameliorated the reduction in cell viability induced by H_2_O_2_-induced stress, the precise mechanism, whether through increasing the number of viable cells via cell proliferation or inhibiting cell death, remains unclear. It is necessary to investigate the effects of these compounds on cytotoxicity in future research.

Notably, the effective concentration of TND1128 (and other compounds) in these studies was lower than that of β-NMN, suggesting the stronger effect of TND1128. This lower effective concentration appears to be attributable to the hydrophobicity and chemical stability of TND1128 and other compounds. Although the four 5-deazaflavin analogs (TNPP0819, TNAD3028, TNCD2611, and TNCD2618) showed comparable activity, and no definitive structure-activity relationship could be discerned, TND1128 had a stronger effect on cell viability than the other derivatives. This study showed the direct effects on cell survival *in vitro*. Given that these derivatives were synthesized with TND1128 as the lead compound to increase brain transfer, it is necessary to examine their effects, including absorption and brain transfer, using animal models of diseases in future investigations.

The Keap1-Nrf2 signaling pathway plays an important role in regulating biological responses to oxidative stress. Nuclear factor erythroid 2-related factor 2 (Nrf2), a member of the Cap’n’Collar (CNC)-BZIP transcription factor family (19), typically forms complexes with Kelch-like epichlorohydrin-related proteins (Keap1) under normal physiological conditions, maintaining a state of low activity. However, under oxidative stress or other pathological stimuli, Nrf2 is released from the complex and translocates to the nucleus. The Nrf2-mediated transcription process is initiated, regulating downstream gene expression, activating a cascade of antioxidant enzymes and phase II antioxidant enzymes, eliminating ROS and other harmful substances, and promoting antioxidative stress, anti-inflammatory, anti-apoptotic, and other cellular protection mechanisms (20, 21, 22).

Various compounds or precursors that activate this pathway, such as sulforaphane in broccoli sprouts and curcumin in turmeric, are present in plant metabolites (23, 24, 25). Furthermore, given that β-NMN attenuates brain injury by activating the Keap1-Nrf2 signaling pathway (26), TND1128 and other compounds may have also exhibited cytoprotective effects via this pathway, similar to β-NMN. In our study, we found that TND1128 and other 5-deazaflavins derivatives, which increase intracellular ATP production through mitochondrial activity, have cytoproliferative and cytoprotective effects. Notably, extracts derived from various fruits and vegetables exhibit neuroprotective properties in both cell culture and animal models, relevant to the pathogenesis of various neurodegenerative conditions, including stroke, Alzheimer’s disease, and Parkinson’s disease (27). Our compounds may also be applicable to numerous neurodegenerative diseases. However, the detailed mechanisms underlying the protective actions of TND1128 and related compounds against oxidative stress, as well as their ability to enhance ATP production, remain unclear and warrant further investigation.

## Data Availability

The data that support the findings of this study are available from the corresponding author upon reasonable request.

## Authors’ Contributions

KK designed the study, performed the experiments, analyzed the data and wrote the manuscript. SK supervised the experiments and revised the manuscript. TW, NA, TN and KI supervised the experiments. All authors read and approved the submitted manuscript.

## Acknowledgments

This study was supported by research grants to K.K. (JSPS KAKENHI, 23K06204) and S.K. (JSPS KAKENHI, 23K27496), and by research grants from Kumamoto Kinoh Hospital and Chemiteras, Inc. to S.K. (Nos. 180231 and 180538).

